# Label-free, real-time monitoring of membrane binding events at zeptomolar concentrations using frequency-locked optical microresonators

**DOI:** 10.1101/2023.09.20.558657

**Authors:** Adley Gin, Phuong-Diem Nguyen, Jeffrey E. Melzer, Cheng Li, Hannah Strzelinski, Stephen B. Liggett, Judith Su

**Author notes:** These authors contributed equally to this work (Adley Gin, Phuong-Diem Nguyen).

## Abstract

Binding events to elements of the cell membrane act as receptors which regulate cellular function and communication and are the targets of many small molecule drug discovery efforts for agonists and antagonists. Conventional techniques to probe these interactions generally require labels and large amounts of receptor to achieve satisfactory sensitivity. Whispering gallery mode microtoroid optical resonators have demonstrated sensitivity to detect single-molecule binding events. Here, we demonstrate the use of frequency-locked optical microtoroids for characterization of membrane interactions *in vitro* at zeptomolar concentrations using a supported biomimetic membrane. Arrays of microtoroids were produced using photolithography and subsequently modified with a biomimetic membrane, providing high quality (Q) factors (>10^6^) in aqueous environments. Fluorescent recovery after photobleaching (FRAP) experiments confirmed the retained fluidity of the microtoroid supported-lipid membrane with a diffusion coefficient of 3.38 ± 0.26 μm^2^ ·s^-1^. Utilizing this frequency-locked membrane-on-a-chip model combined with auto-balanced detection and non-linear post-processing techniques, we demonstrate zeptomolar detection levels The binding of Cholera Toxin B-monosialotetrahexosyl ganglioside (GM1) was monitored in real-time, with an apparent equilibrium dissociation constant (k_d_) = 1.53 nM. The measured affiny of the agonist dynorphin A 1-13 to the κ-opioid receptor revealed a k_d_ = 3.1 nM using the same approach. Radioligand binding competition with dynorphin A 1-13 revealed a K_d_ in agreement (1.1 nM) with the unlabeled method. The biosensing platform reported herein provides a highly sensitive real-time characterization of membrane embedded protein binding kinetics, that is rapid and label-free, for toxin screening and drug discovery, among other applications.

## INTRODUCTION

The cell membrane is comprised of lipids and transmembrane proteins and is fundamental for controlling the signal transduction events essential for cellular communication and regulatory responses. As much as 30% of the cell interface is estimated to be integral membrane proteins,^1^ which represent a wide variety of functions. This includes G protein coupled receptors (GPCRs, currently the most common therapeutic target), various channels and pumps, immunoglobins, and virulence receptors. In addition, lipids within the membrane act as receptors as well, such that the cell membrane serves as a key interface for recognizing and responding to extracellular phenomena. The lipid composition of the membrane itself can also affect the cellular consequences of activation or blockade, due to its influence on transmembrane protein conformation.^2^ While a significant portion of pharmaceutical compounds are designed to target membrane components such as receptors, they are often difficult to label. Radioisotopes can break bonds and induce conformational changes and fluorescent labels are large in comparison to small molecules and can alter their chemical behaviors.^3^ Typical screening methods are also limited by the ability to control the membrane environment of the protein. The ability to observe membrane receptor-drug interactions in real-time without the use of labels could enhance the drug discovery process as well as increase our understanding of physiological events.

Conventional methods for studying membrane-protein binding events include the use of *in vivo* animal models^4^ (typically functional readouts with implied affinities) or selected cells *in vitro*^5^. Enzyme linked-immunosorbent assays (ELISA),^6^ liposome microarrays,^7,8^ radioisotope labeling assays,^9^ and fluorescent imaging^10^ are also methods used with cells or tissue homogenates to ascertain these interactions, each having limitations such as those described above. Of particular interest are label-free sensing schemes, which can directly and rapidly detect the binding of molecular ligands to receptors targets. Several examples that can quantify binding kinetics are surface plasmon resonance^11^ (SPR) with a limit of detection (LOD) down to 1 pg/mm^2^, electrochemical biosensors,^12,13^ and electrochemical impedance spectroscopy (EIS) (LOD < 10 pM).^14^ Complex and potentially interfering signal enhancement tags are often required to boost the sensitivity of these systems.^15–17^

Whispering gallery mode (WGM) optical biosensors are ultra-sensitive, label-free, and capable of observing single molecule binding events. They confine photons in a path circumscribing the cavity; the photons can circulate many times thus allowing small changes in the optical path, such as those caused by biomolecular binding.^18–23^ Here we measure lipid membrane formation and membrane binding events by utilizing an ultra-sensitive, label-free biosensing system known as FLOWER (frequency locked optical whispering evanescent resonator).^18,19,24–27^ FLOWER is able to detect a single macromolecule; the high quality (Q)-factor and the evanescent field of WGM microtoroid resonators are exploited so that any local refractive index changes caused by analyte binding events to the resonator’s surface can be measured in real-time as a shift in the cavity’s resonance frequency (Fig. 1).^24,25,28,29^ Previous applications of WGM microresonators include detecting performance enhancing drugs^28^, nucleic acids^30–32^, proteins^25,32– 34^, volatile organic compound sensing,^26^ protein interaction screening^35,36^, and frequency comb generation.^37,38^ Although previous studies have demonstrated the use of optical microcavities for studying the formation of lipid membranes^39^ and protein-lipid interactions^35^, to the best of our knowledge, optical microcavities have not been used to observe membrane receptor-ligand binding kinetics and associated affinity measurements. The ability to detect these interactions in a real-time, label-free, and ultrasensitive manner will significantly advance biomarker screening as well as drug discovery, among other applications.

**Fig. 1.**
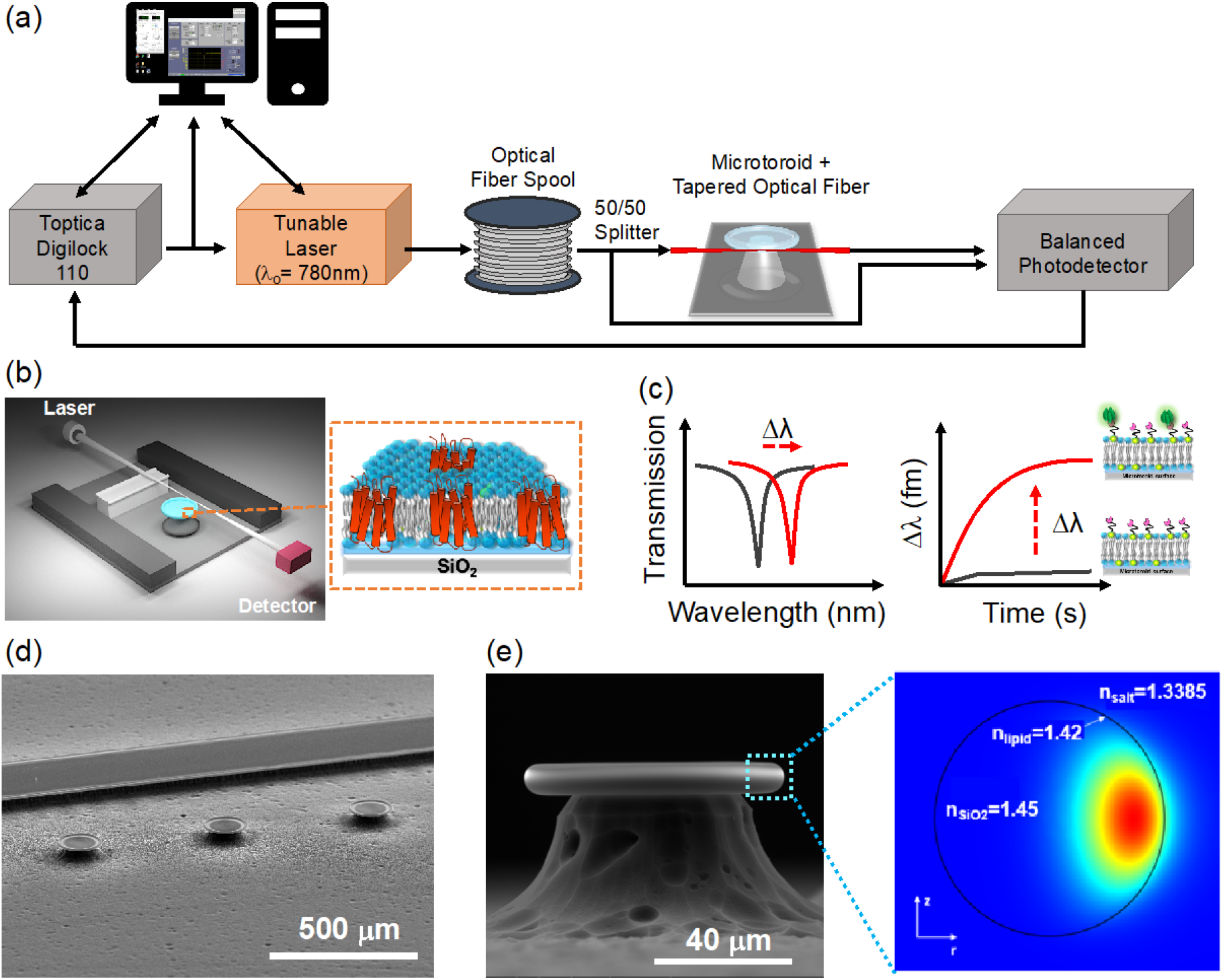
FLOWER system. (a) A tunable laser is coupled to the microtoroid cavity through a tapered optical fiber. (b) Schematic diagram of the constructed biosensing chamber (top cover glass not shown). The tapered fiber is glued to the support wall with a polymer adhesive. Inset shows the structure of a membrane protein (red) incorporated in a lipid bilayer (blue) on the toroid’s glass surface. (c) Sketches of how the resonant wavelength red-shifts when analytes adsorb onto the functionalized toroid surface. (d) SEM image of microtoroid array alongside a support wall. (e) SEM sideview image of a microtoroid structure. The inset picture shows a COMSOL simulation of an optical mode of the lipid coated cavity in a buffered aqueous solution.

In our approach, we establish a methodology for directly investigating membrane binding events on a WGM microtoroid resonator (Fig. 1) using two different ligand-receptor schemes: Cholera Toxin B (CTB) binding to monosialotetrahexosyl ganglioside (GM1) membrane receptors and Dynorphin A 1-13 (Dyn A 1-13) binding to the κ-opioid GPCR (kOR). Respectively, these represent a pathologic signaling cascade leading to the symptoms of cholera and a neuronal target for treating mood and other stress reponses^40,41^ To mimic a native lipid environment for studying membrane binding events, a synthetic lipid bilayer was first self-assembled by rupturing lipid vesicles onto the microtoroid’s dielectric silica surface. The GM1 glycolipid receptors were prepared together with the lipid membrane,^8^ while κ-opioid receptors (κOR) were incorporated by means of micelle dilution.^42^ The binding kinetics of CTB to GM1 and Dynorphin A 1-13 to κOR, respectively, were measured without labels, in real-time (i.e. minutes) with minimal (30 µl) sample consumption and low LOD (150 zM for the Dynorphin A 1-13 assay) offering a significant advantage over other molecular assays. To confirm the specificity of the binding assay, fluorescent imaging and radioligand competitive binding assays were performed. Our results demonstrate a platform for rapid, label-free, ultra-sensitive measurements of lipid biomembrane formation, small molecule binding, and protein binding kinetics, among other applications.

## RESULTS

### FLOWER biosensing system

Fig. 1a and 1b illustrate the approach, including the microtoroid functionalized with a proteolipid membrane. The tuning range of the laser used in these studies was 765 nm to 781 nm, conditions where light absorption in water is minimal. Light from the laser is evanescently coupled into the toroid using a tapered optical fiber, although free-space coupling of light into the toroid can be done as well.^43,44^ The tunable laser is frequency-locked to the microtoroid resonator so that the laser wavelength matches the microtoroid resonance. To calculate the Q-factor of a resonance, the spectral dip was fitted to a Lorentzian curve and the resonant wavelength was divided by the full-width-half-max (FWHM).^39^ The dip in the transmitted intensity occurs when the optical path length of the resonator is equal to an integer multiple of the circulating light’s wavelength:

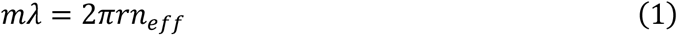

where *m* is an integer representing the number of wavelengths per round trip, λ is the free-space wavelength of the laser, *r* is microtoroid major radius, and *n*_*eff*_ is the effective refractive index (RI) of the guided mode. As protein or lipid materials land on the microtoroid surface within the evanescent light field, the effective refractive index of the optical mode increases, causing a shift in resonance frequency proportional to the number and size of analytes bound (Fig. 1c).^25,30,45^ A two-dimensional axisymmetric COMSOL simulation was performed to aid in visualizing the effective sensing region at the equator of the lipid coated microtoroid (Fig. 1 d,e) (See Methods for details). By choosing RIs in our simulations comparable to experimental values for the microtoroid cavity, lipid materials, and surrounding aqueous solution, we can see that a portion of the optical mode extends past the surface and into the lipid layer and aqueous environment. FLOWER has an advantage over plasmonic sensors^46^ in that the toroid has a larger capture area thus enabling faster detection times.^29^

### GM1-DOPC lipid functionalization of microtoroid optical resonators

A synthetic phospholipid membrane composed of DOPC lipids doped with 2% mol GM1 receptors was used to functionalize the silica microtoroids for quantifying CTB binding affinity (Fig. 2a). DOPC is a zwitterionic lipid membrane that can be used as an antifouling membrane to resist non-specific binding of proteins to silica (Fig. S1 & S2). Unilamellar GM1-DOPC lipid vesicles were produced by extruding the GM1-DOPC lipid suspension through a 100 nm pore filter (see Methods) and their size was confirmed using dynamic light scattering (Fig. S3a). A GM1-DOPC lipid bilayer can be quickly formed on the silica toroid surface as the hydrophilic head of the lipid vesicles readily rupture and adsorb onto the toroid (Fig. 2b). Membrane fluidity on the toroid surface was measured by fluorescent recovery after photobleaching (FRAP) (Fig. S3b), which shows the lipid bilayer retaining 80% membrane fluidity with a diffusion coefficient, *D*, of 3.38 ± 0.26 μm^2^ s^-1^ which is in the range typically observed for supported lipid bilayers formed on glass^57,58,59^.

**Fig. 2.**
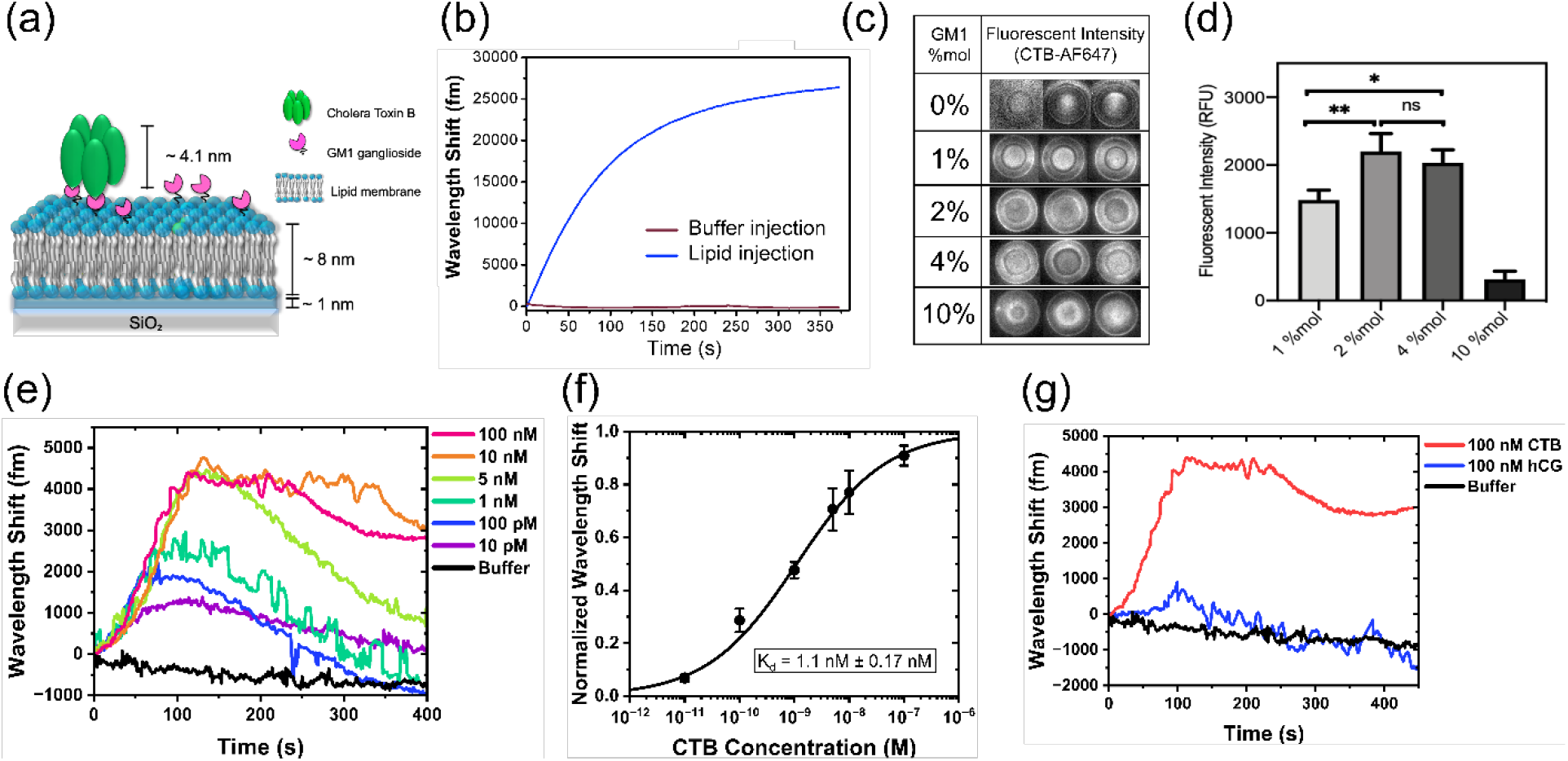
GM1-CTB binding signals. (a) Schematic model of pentameric CTB bound to a GM1-DOPC lipid membrane silica microtoroid. (b) Wavelength shift from GM1-DOPC lipid vesicles adsorbing onto the silica toroid. (c) Fluorescent images of 50 nM CTB-AF647 binding to varying % mol fraction GM1 in DOPC coated-microtoroids. (d) Fluorescent intensity determined from the fluorescent assay in c). Data are mean ± SD (n = 6-9 experiments). Statistical significance was determined using a one-way ANOVA, \**p* = 0.028, \*\**p* = 0.006, *ns*: not significant). (e) Wavelength shift as CTB binds to a GM1-DOPC functionalized toroid. (f) Dose-response binding curve generated based on the maximum wavelength shift from e). Data are mean ± SEM from 3 independent experiments. (g) Control experiment attempting to detect an irrelevant protein (hCG) binding to the GM1-DOPC coated microtoroid.

For bare microtoroids, we generally obtained Q-factors in the range of 10^6^-10^7^ in aqueous buffer solution. The Q-factors of optical microresonator cavities in aqueous solutions are typically lower than in air due to the smaller difference in refractive index between the resonator material and surrounding environment (Fig. S3c). After lipid functionalization, microtoroids still maintained a high Q-factor in the range of ~10^6^ which is suitable for high-sensitivity biosensing (Fig. S3d).

### FLOWER detects CTB binding

Cholera toxin B is a subunit of the cholera toxin secreted by the bacterium *Vibrio cholera* and binds to GM1 glycolipid receptors presented on the surface of cells. The GM1 receptor is responsible for the binding and internalization of this virulent protein, resulting in the subsequent rapid loss of fluids from the intestine and severe diarrhea presented in cholera.^12,47^

CTB binding to a GM1-DOPC functionalized toroid was demonstrated *in vitro* by sequentially injecting increasing concentrations of unlabeled CTB prepared in buffer into a fluidic chamber containing the microtoroid (Fig. S8) and tracking the wavelength shift as done previously with our FLOWER system (Fig. 1a).^24,25,28^ The shape and kinetics of the WGM response was consistent, exhibiting a rapid and positive wavelength shift and peak at t~120s after the start of the sample injection, followed by a gradual decline back to the baseline for CTB concentrations <10 nM (Fig. 2e). The WGM response exhibited a net positive wavelength shift at steady state for higher CTB concentrations (>10 nM). It should be noted that CTB is being continuously injected (~1.2 ul/s) when the WGM response is being recorded in Fig. 2e (t=0s to t=400s). Between each CTB injection, buffer is injected for 10 minutes to rinse the fluidic chamber and bring the sensor back to a steady state (data not shown). As expected, the peak of the WGM response increased along with the CTB concentration (Fig. 2e). The shape of our WGM response is similar to the dynamic mass redistribution (DMR) response from resonant waveguide grating biosensors, which also rely on the evanescent field for label-free detection.^48–51^

The binding curve was generated using the peak of the WGM response for each CTB concentration and fitted using the Hill-Waud binding model (Fig. 2f).^52,53^ The dissociation constant, K_d_, of the GM1-CTB binding interaction was 1.1 nM. For our particular system, which uses a 2% mol doping density for GM1, our K_d_ value of 1.1 nM is in reasonable agreement with the K_d_ value of 0.73 nM that was obtained via surface plasmon resonance for the same doping density.^54^ Surprisingly, the Hill coefficient of the fit was *n* = 0.52, suggesting some level of negative binding cooperativity between the CTB and GM1 receptor, while previous literature reported positive cooperativity (i.e. n>1).^52,55^ However, the previously reported experiments were performed using supported lipid bilayers on flat surfaces, while the curvature of the toroid geometry at the sensing region may decrease the cooperative effect between the CTB and GM1 receptor.

### Determination of optimum concentration of GM1 in the lipid membrane

A high concentration of GM1 in the lipid membrane results in clustering of the receptor or other effects, and decreased CTB binding efficiency.^55,56^ Thus, we investigated the optimum concentration of GM1 doping on the lipid coated microtoroid. A fluorescent assay was performed using multiple microtoroid substrates doped with varying GM1 doping ratios (0 to 10 % mol). For each substrate, 50 nM of CTB labeled with Alexa Fluor 647 (CTB-AF647) was incubated and the binding was measured using fluorescent imaging (Fig. 2c, 2d). The amount of doped GM1 glycolipid in the membrane has a remarkable effect on the association of CTB molecules. We observed that at 10 % mol GM1 the fluorescent signal on the toroidal microcavity was significantly decreased, whereas a maximum in the fluorescent signal was observed with 2 and 4 % mol of doped GM1. This phenomenon is in agreement with existing literature.^55,56^ Alternatively, the weakened binding of CTB to GM1 can also be explained by the electrostatic repulsion between negatively charged CTB and increasing negative charge of lipid membrane by increased GM1 ligand density (each GM1 headgroup has (-1) charge).^55^ The specificity of the detection was confirmed by introducing an irrelevant protein (HCG, human chorionic gonadotropin) onto a GM1-functionalized microtoroid (Fig. 2g). hCG has very low affinity for the GM1 receptor.^57^ The injection of 100 nM hCG produced a red-shift near the baseline noise (not statistically different than buffer) which was several thousand fold lower than the CTB response.

### FLOWER detects DynA 1-13 - κOR interactions

κ-opioid receptors (κOR), together with μ- and σ-opioid receptors are an important mediator system for emotional and behavioral responses to stress and pain. The binding of the endogenous ligand dynorphin during stress activates κOR to produce analgesia and dysphoria.^58^ The dysregulation of central nervous system responses to stress can elevate anxiety, which can lead to depression and drug seeking behavior.^58–60^ Here, we explore the potential use of our lipid membrane coated optical microcavity for drug screening purposes, demonstrated by a κOR proteolipid functionalized microtoroid sensor for the detection of unlabeled DynA 1-13. The proteolipid membrane formation used in this study is shown in Fig. 3. The microtoroids were incubated in DOPC lipid (0.5 mg/ml) to form an artificial lipid bilayer on the silica sensing surface of the sensor. The formation of a lipid bilayer on the microtoroid was confirmed by measuring the resonant wavelength shift as DOPC lipid was injected into the fluidic chamber; at t=1750s buffer is injected to wash away excess lipid vesicles (Fig. 4a). HEK293T cells were transfected with the cDNA encoding the human κOR, with expression verified by immunoblots of lysed cells and intact cell confocal imaging (Fig. S9). κOR-expressing cells were solubilized using the detergent CHAPS in a lysis lysis buffer to form κOR micelles prior to sensor deposition (see Methods). The κOR micelles were incorporated into the lipid bilayer on the microtoroid by means of micelle dilution.^61^ Successful incorporation of κOR was confirmed by measuring the resonant wavelength shift (Fig. 4b) and by fluorescence imaging (Fig. S4). Note that a relatively low concentration of κOR was used in Fig. 4b, yet a large spectral shift was produced immediately following receptor addition, suggesting a strong incorporation efficiency.

**Fig. 3.**
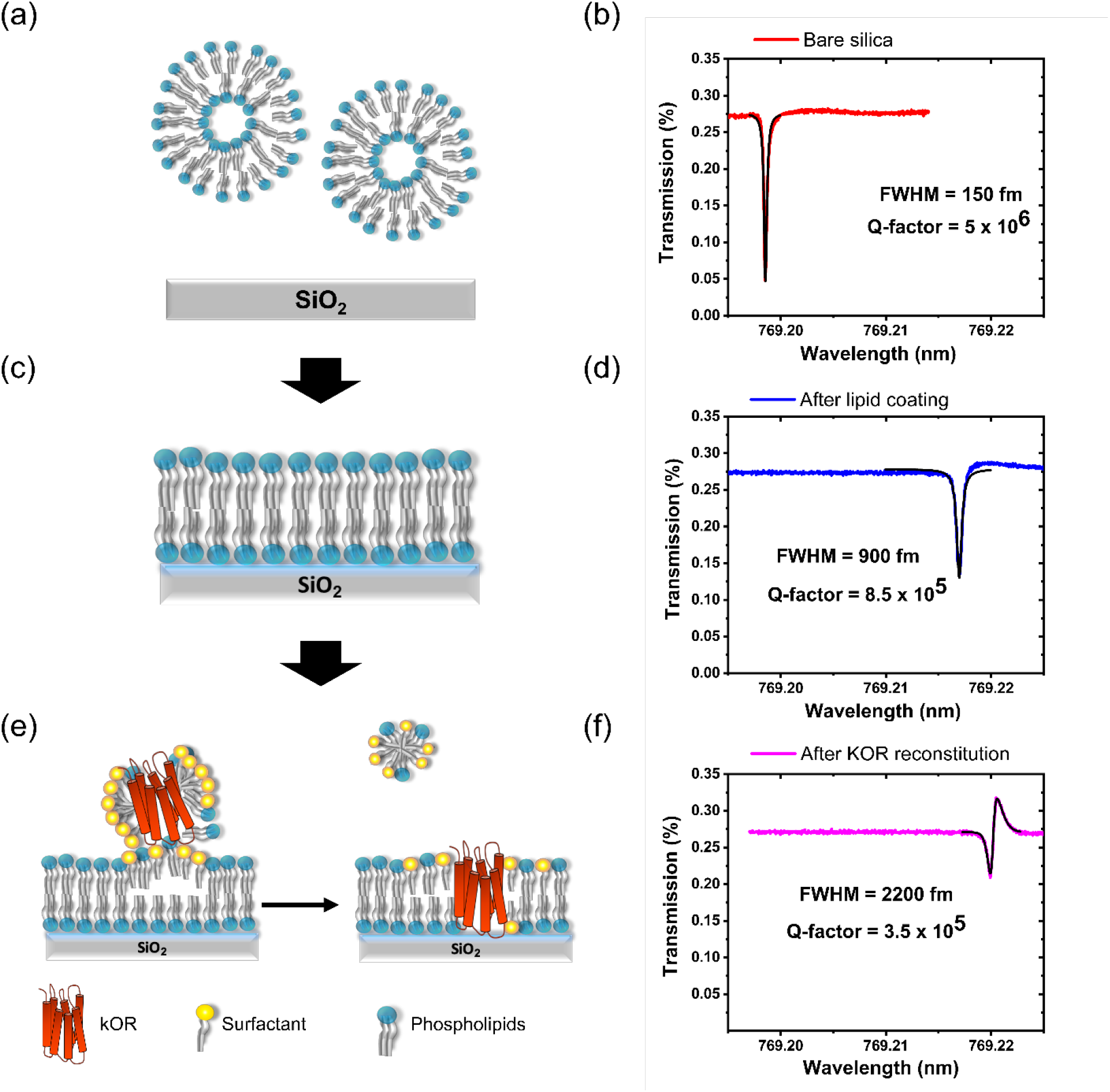
Resonant wavelength shift quantification of κOR proteolipid membrane formation on a microtoroid optical resonator for DynA 1-13 label-free detection. (a,b) Just before lipid adsorption, (c,d) after lipid coating process, (e,f) after κOR reconstitution by means of micelle dilution.

**Fig. 4.**
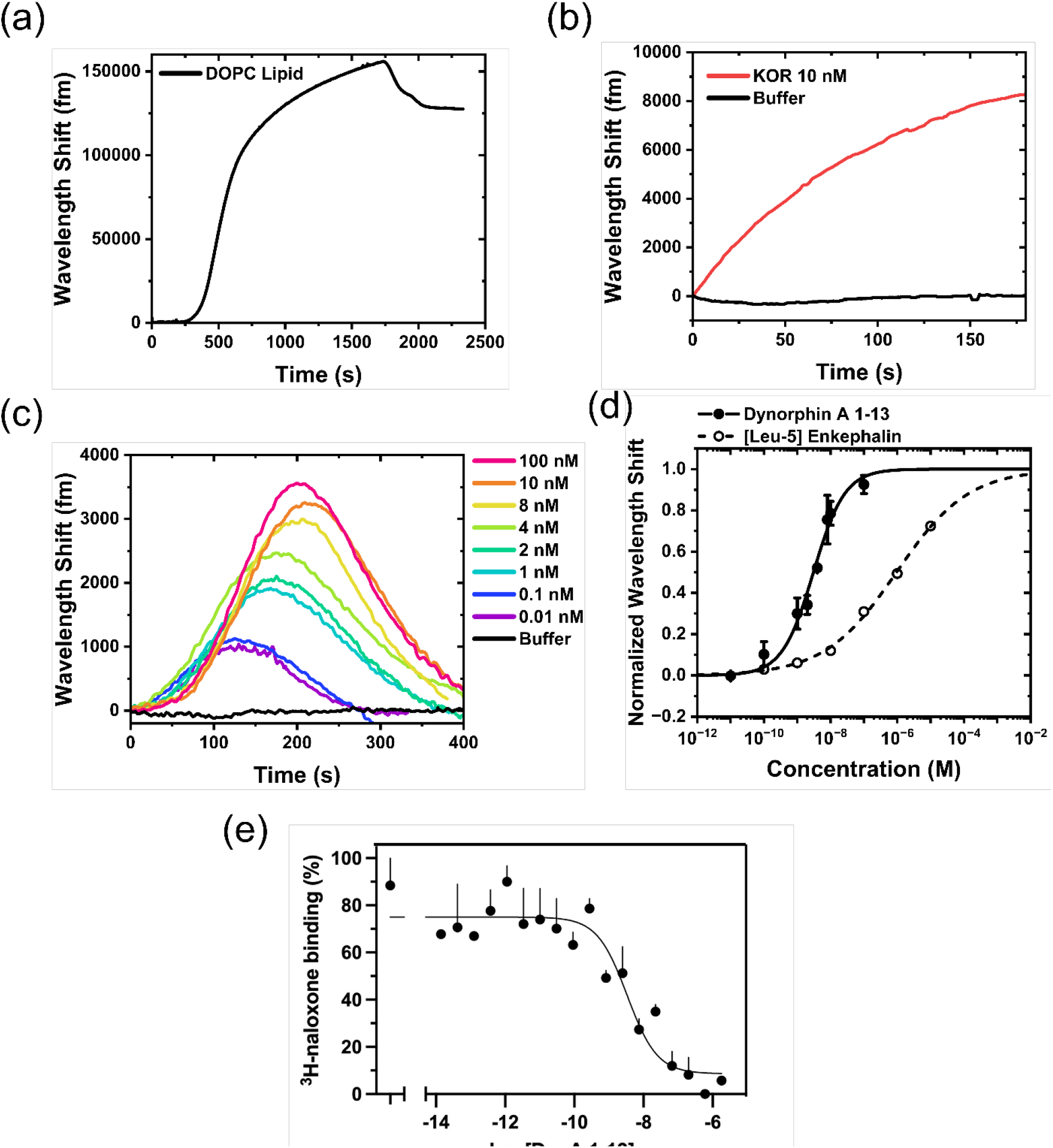
Quantification of κOR/DynA 1-13 binding kinetics using a proteolipid-coated microtoroid optical sensor. (a) WGM shift as lipid vesicles adsorb onto a bare silica toroid. (b) WGM shift as κOR is incorporated into the lipid bilayer on the toroid. (c) WGM shifts as a function of increasing concentrations of unlabeled DynA 1-13 binding to κOR-lipid toroid sensor. (d) Binding curve of DynA 1-13 (solid line) and [Leu5]-enkephalin (dashed line) to κOR-lipid toroid. Error bars represent the standard error of three independent measurements. (e) DynA 1-13 competes for binding of the κOR radioligand 3H-naloxone in the membranes from κOR-transfected cells. Results are mean ± SEM from 3 experiments.

DynA 1-13 is a tridecapeptide, a major metabolite from the endogenous Dynorphin A.^62^ The study of DynA 1-13 binding kinetics has been significantly challenging due to its inaccessibility to labeling techniques. Here, the assay was performed by injecting increasing concentrations of unlabeled DynA 1-13 into the experimental fluidic chamber containing the κOR-lipid functionalized toroid (Fig. S8). The WGM response consistently exhibited a rapid red-shift and peak (t~100-200s) after the initial injection of the κOR sample followed by a decline back to the baseline (Fig. 4c). Note that DynA was continuously injected for the entire duration of recording the WGM response (t=0 to 400s). Buffer was injected for 10 minutes between each DynA injection to rinse the chamber and bring the sensor back to a steady state. The peaks of the WGM shifts were fitted to the Hill-Waud specific binding model yielding a K_d_ of 3.1 ± 0.3 nM (Fig. 4d). Depending on the experimental methods, the K_d_ has variously been reported to range from 0.25 nM - 6.1 nM for DynA1-13 at the κOR.^60,63,64^ The estimated LOD for the label-free DynA 1-13 detection assay was 150 zM. A fluorescent imaging assay was conducted to ascertain nonspecific interactions. We conducted the fluorescent assay experiment using biotinylated DynA 1-17 and streptavidin labeled AF647. Fig. S4 shows the fluorescent imaging results of DynA 1-17 binding to the κOR reconstituted lipid coated microtoroid. In the absence of κOR in the lipid membrane, changes in the fluorescent intensity were insignificant. This data suggests that the κOR binding function to DynA 1-13 molecule is retained using the micelle dilution method, and the observed binding kinetics of κOR receptors characterized by our microtoroid sensing system is specific to the DynA 1-13 molecule. To demonstrate rank-order specificity of the κOR toroid sensor, the experiment was repeated with the low-affinity κOR agonist [Leu5]-enkephalin. [Leu5]-enkephalin is another endogenous opioid peptide which is known to have agonist action at both the μ- and δ-opioid receptors, but weakly binds to the κOR subtype (compared to DynA 1-13).^60,65^ The K_d_ of [Leu5]-enkephalin at the κOR was determined experimentally to be 947 ± 92 nM (Fig. 4d).

To confirm our DynA 1-13 affinity determination in a parallel manner, we performed radioligand binding assays using the same recombinantly expressed k-OR in HEK-293T cells as was used in the microtoroid studies. Here, ^3^H-naloxone was used as a label for the receptor. Cell membranes were incubated with 5 nM of this radiolabel and varying concentrations of DynA 1-13, and the bound ^3^H-naloxone captured by filtration and washing over glass fiber filters (see Methods). The K_d_ for DynA 1-13 derived from these competition studies was 1.1 ± 0.84 nM (Fig. 4e), in good agreement with the WGM data.

In a separate experiment for determining the LOD, DynA 1-13 was injected while the wavelength shift was measured (Fig. 5a). For each DynA 1-13 concentration, the peak wavelength shift was taken and plotted on a calibration curve. The mean, 1SD, and 3SD values from repeated buffer (i.e. blank) injections are shown as a horizontal line and shaded areas in the calibration curve (Fig. 5b). The LOD was defined as the minimum detectable DynA 1-13 concentration above the background signal mean + 3SD, which was calculated to be 150 zM.

**Fig. 5.**
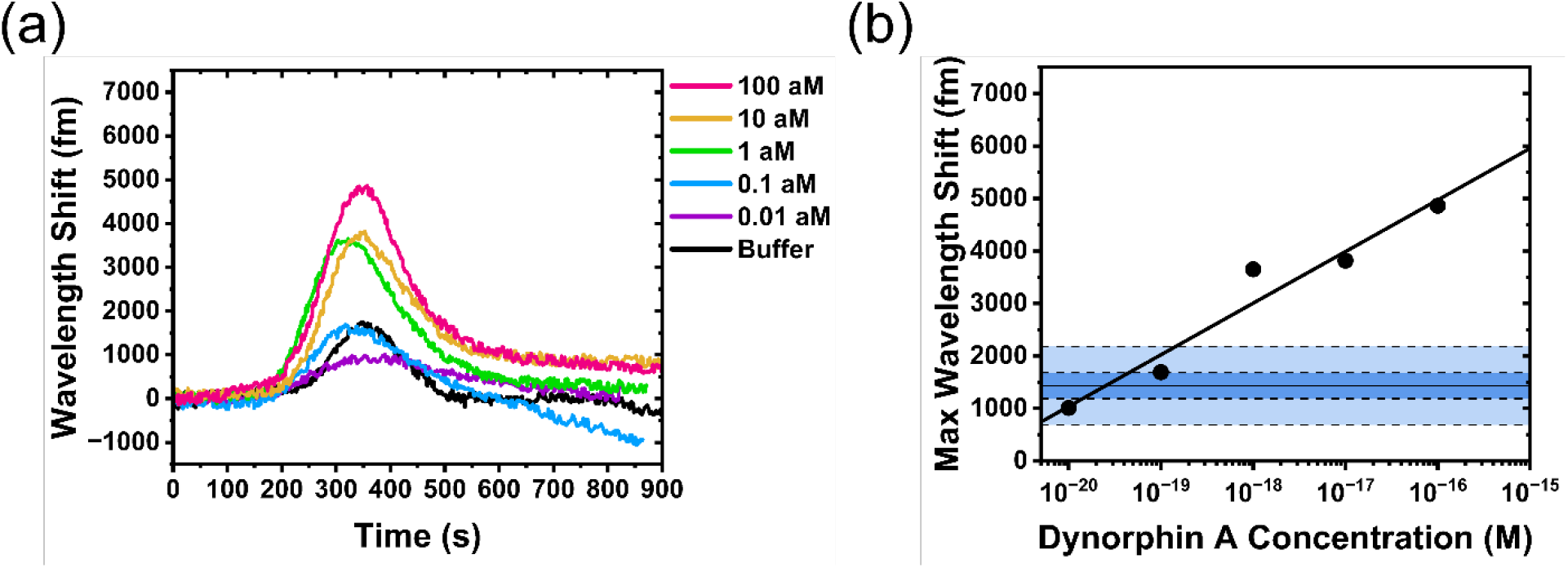
(a) Separate experiment showing attomolar DynA 1-13 binding to the κOR-lipid toroid. (b) Calibration response curve for attomolar DynA 1-13 binding experiment. Shaded areas show ±1SD and ±3SD from the mean of repeated buffer injections.

### Two-drug competitive binding experiments with kOR functionalized-microtoroid

Another way to assess drug targeting to receptors is to bind a drug to the receptor and then ascertain the output effects of increasing concentrations of a second ligand (similar to the radioligand competition experiment in Fig. 4e). Using FLOWER, we considered that the wavelength shift from bound DynA 1-13 would decrease with increasing concentrations of the opiate antagonist naloxone, indicating its applicability in this type of pharmacologic assay. To test this, we varied naloxone concentrations from 100 pM to 1 μM while keeping the DynA 1-13 concentration constant (10 nM). The increasing concentrations of naloxone indeed competed with the binding of DynA 1-13 to κOR, resulting in the decreases in the peak resonant wavelength shifts as the naloxone concentration increased (Fig. 6a). A high and low IC_50_ was obtained by fitting the competitive response curve with a two-site competitive binding model (Fig. 6b). The K_i_ was calculated from the IC_50_ using the Cheng-Prusoff equation (Eqn. 5). The low K_i_ was calculated to be 92 nM and the high K_i_ was calculated to be 0.26 nM. The basis of this 2-site fit is not altogether clear, but this has also been observed using radioligand binding with similar affinity ratios.^66^ .

**Fig. 6.**
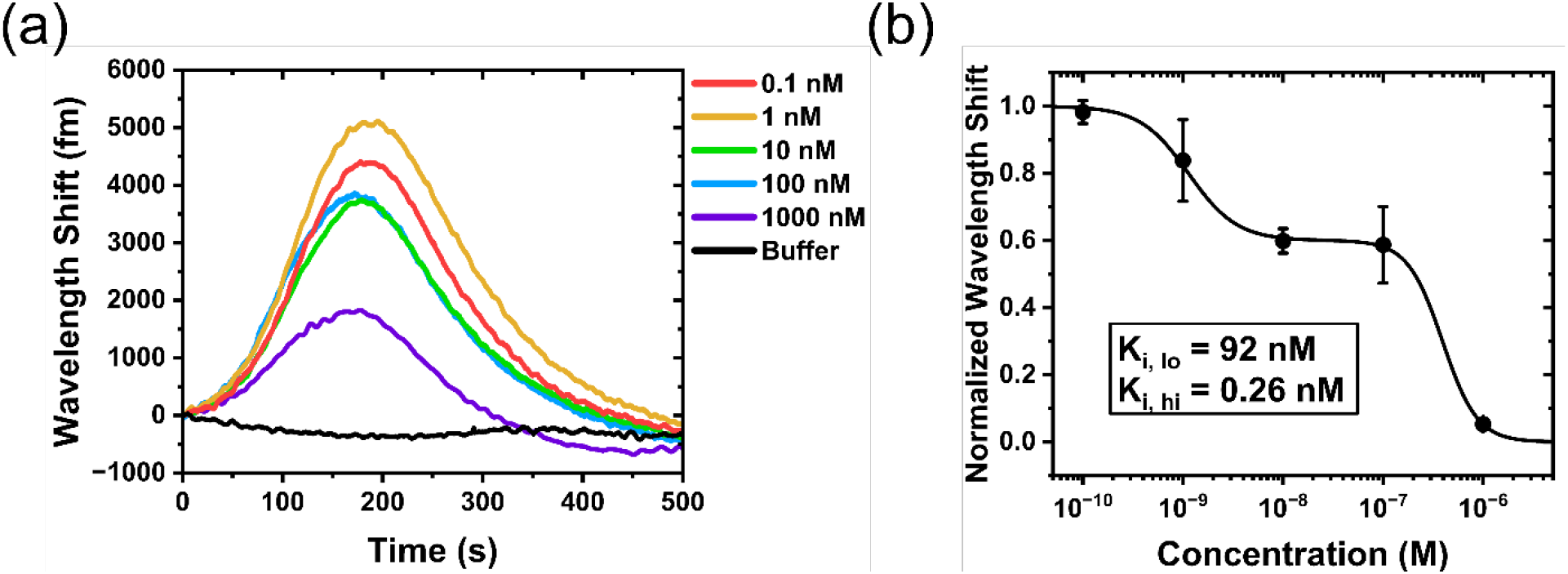
(a) WGM shift showing the competitive binding of naloxone with 10 nM DynA 1-13 to κOR binding sites. (b) Competitive binding curve of naloxone vs. 10 nM DynA 1-13 on a κOR-lipid toroid.

## CONCLUSION

FLOWER offers real-time, rapid characterization of biomimetic membrane formation, protein incorporation, and measurement of lipid membrane receptor-ligand binding kinetics and associated affinity measurements without the use of labels. An *in vitro* biomimetic membrane model can be formed on the dielectric microtoroid surface to serve as a sensitive detection interface for direct tracking of lipid fusion and membrane protein incorporation, as well as subsequent ligand binding events. Fluorescent imaging confirmed that the activity of reconstituted membrane proteins was preserved together with the respective ligand recognition ability. Without the complexity of a labeling tag, unlabeled ligands were directly detected resulting in a rapid sample-to-result diagnosis within several minutes. High affinity binding of CTB molecules to GM1 receptors, as well as the study of endogenous opioid peptide neurotransmitters such as Dynorphin A 1-13, [Leu5]-enkephalin and the opioid antagonist naloxone to the κ-opioid receptor were successfully characterized in a label-free, real-time manner. The limit of detection (zeptomolar) obtained in this study is several orders of magnitude lower than reported with other label-free technologies such as SPR, electrochemical impedance spectroscopy (EIS), and microring resonators. These technologies have a detection range usually in the nM to pM level. The present study not only demonstrates the promising potential of using FLOWER for drug and toxin screening research, but also provides a powerful platform for advancing knowledge in membrane interactions *in vitro*, suggesting a new approach for preliminary therapy and disease prevention.

## MATERIALS AND METHODS

### Materials

The silicon substrate was purchased from University Wafer. The buffered oxide etchant (6:1 v/v, KMG, Texas, USA), Tris-HCl, ethylenediaminetetraacetic acid (EDTA), isopropanol, methanol, chloroform, Triton X-100, dimethyl sulfoxide (DMSO), Trizma®, unlabeled Cholera toxin B subunits (CTB), Dynorphin A 1-13 (YGGFLRRIRPKLK), and naloxone were all purchased from Sigma Aldrich (St. Louis, MO). Avanti Polar Lipids (Alabaster, AL, USA) supplied all the lipids used in the experiments: 1,2-dioleoyl-sn-glycero-3-phosphocholine (DOPC); Ganglioside GM1 from ovine brain (GM1 ganglioside); 1,2-dioleoyl-sn-glycero-3-phosphoethanolamine-N-(Lissamine rhodamine B sulfonyl) (ammonium salt) (LissRhod:PE); 1,2-distearoyl-sn-glycero-3-phosphoethanolamine-N-[poly(ethylene glycol)2000-N’-carboxyfluorescein] (ammonium salt) (PEG2000-PE-CF); 1,2-distearoyl-sn-glycero-3-phosphoethanolamine-N-[amino(polyethylene glycol)-2000] (ammonium salt) (DSPE-PEG2000-Amine) . Cholera Toxin B subunit labeled with Alexa Fluor 647 (AF647), and Alexa Fluor™ 488 NHS Ester (Succinimidyl Ester) were purchased from Thermo Fisher (Watham, MA). [Leu5]-enkephalin (YGGFL), biotinylated Dynorphin A 1-17 (Biotin – YGGFLRRIRPKLKWDNQ) were purchased from Anaspec (Fremont, CA). All solutions were prepared with 18.2 MΩ·cm water passed through a 0.2 μm polycarbonate Whatman filter and degassed to remove bubbles before use.

### Microtoroid Fabrication

Microtoroid structures were fabricated as discussed previously.^19,40,41^ In brief, fabrication of a microtoroid started with a 2 µm thick layer of SiO_2_ on a Si wafer (universitywafer.com). Disks with diameter 150 µm along with a 350 µm wide support wall were patterned onto the SiO_2_ surface with Microposit S1813 photoresist. Afterwards, buffered oxide etchant (BOE) 6:1 was used to etch the exposed SiO_2_ down to the Si substrate, followed by resist stripping with acetone and isopropanol. After a dehydration bake at 130°C, a XeF_2_ dry chemical etcher (Xactix e2, Orbotech, Yavne, Israel) was used to etch the exposed Si substrate. The microdisks and support wall were then reflowed using a CO_2_ laser (Synrad 48-1, WA, USA) to create SiO_2_ microtoroids with major diameter around 100 µm on a 50 µm Si pillar.

### Lipid Preparation

Lipid vesicles were prepared according to a well-established protocol.^42,43^ For GM1-CTB studies, a controlled mol% of methanolic GM1 ganglioside was mixed with DOPC lipid and dried under argon gas. For κOR pharmacokinetic studies, DOPC lipid vesicles were prepared using a 100 nm membrane filter and mini extruder kit from Avanti Lipids. DOPC lipid vesicles were prepared and then stored at 1 mg/ml in PBS at 4°C. Fluorescent doped lipid vesicles were prepared by adding 1% mol of either Liss-RhodPE or PE-PEG2000-CF into GM1/DOPC lipid mixture with (Fig. S2). The average size of the lipid vesicles was characterized by dynamic light scattering (DLS) and transmission electron microscopy (TEM). Lipid vesicles were stained using1 mg/ml uranyl acetate and air-dried overnight before TEM imaging.

### Fluorescent Recovery After Photobleaching (FRAP) Assay

To assess the mobility of the lipid membrane on the microtoroid, FRAP experiments were performed with a Leica SP5-II confocal microscope using GM1 doped DOPC labeled with AF488. A 488 nm Argon laser line at 50 mW was used for photobleaching. Images of fluorescent recovery were taken using a 20x/0.75NA objective and the data was analyzed using ImageJ.

### Microtoroid Biosensing Setup

An optical fiber (Thorlabs SM600, Newton, NJ) was tapered using a stationary hydrogen flame and a custom fiber-pulling stage (Newport, CA, USA). Light from a tunable laser (New Focus TLB-6712, Newport, CA, USA) was coupled into the fiber and the transmission was measured using an auto-balanced photodetector (Nirvana 2007, Newport, CA, USA). After surface functionalization, the microtoroid chip was affixed into a 3D-printed fluidic chamber (internal volume ~120 μl) using double sided tape. A glass cover slip was cut to size and placed on top of the fluidic chamber to retain the fluid. Fluid samples were perfused into the chamber using a 16-channel syringe rack and electric rotary valve system (ASP-ERV-O1.2-16, Aurora Pro Scientific). The microtoroid was evanescently coupled with the tapered fiber using a 3-axis micrometer stage and a 3-axis piezo nanopositioning stage (P-611.3 NanoCube, PI, MA).

### GM1-CTB Bioassay

For lipid bilayer formation on the microtoroid, DOPC vesicles (0.5 mg/ml in Tris-HCl 25 mM, 100 nm in diameter) consisting of different mol % of GM1 were introduced onto the microtoroid chip. The vesicle rupture and fusion process was induced immediately by the edge of the hydrophilic silica microtoroid.^44,45^ The lipid coating process was quantified in real-time by tracking the resonance frequency shifts, then washed thoroughly with the binding buffer (Tris-HCl 25 mM, NaCl 150 mM, EDTA 1 mM, pH 7.4). For the CTB binding assay, increasing concentrations of CTB prepared in binding buffer were sequentially introduced into the microtoroid sensing chamber. A control experiment was conducted using fluorescently labeled lipids to confirm lipid stability upon multiple cycles of buffer washing (Fig. S5).

### κOR studies

For FLOWER, the lipid bilayer was formed on the microtoroid by incubating the toroid chip in a DOPC vesicle solution (0.5 mg/ml, 100 mM phosphate buffered saline) for 1 hour at room temperature. Afterwards, the toroid chip was washed with PBS and then transferred to Tris buffer (Tris-HCl 25 mM, NaCl 150 mM, EDTA 1mM, BSA 0.05% w/v, pH 7.5). κOR were generated by transfecting HEK-293T cells with an expression vector consisting of the human kOR coding sequence with an amino-terminal peptide tag (HA) in the expression vector pcDNA3. Transfections were performed using Lipofectamine 2000, and after 2 days the cells were washed with PBS, pelleted and frozen at -80°C.^67^ Non-transfected HEK-293T cells were used as a negative control. Expression in cell membranes was confirmed by immunoblotting using an HA-specific antibody as described.^68^ The same antibody was used for fluorescent confocal microscopy with fixed cells as described.^69^ K-or were isolated from the HEK293T crude membranes using a combination of shearing with a syringe and detergent solubilization .^61^ Briefly, 100 μl of lysis buffer (CHAPS 10 mM, Tris-HCl 25 mM, NaCl 150 mM, EDTA 1 mM, AEBSF 1 mM, pH 7.5) was added to ~80 mg of cell membranes in a 1 ml centrifuge tube. κOR extraction from the membrane was aided by repeated passage of the suspension through a 23-gauge needle attached to a 1 ml syringe. The cell suspension was incubated on ice for 1 hour and then centrifuged for 30 minutes at 15,000 x *g* at 4°C. The resulting supernatant with the κOR-detergent micelles, was reconstituted into the lipid coated microtoroid by the detergent dilution method.^42,61^ In brief, 100 ul of the KOR supernatant was added to a 1 ml centrifuge tube containing 100 ul of Tris buffer and the lipid coated toroid chip. Detergent dilution below the CHAPS CMC (8-10 mM) results in spontaneous insertion of the κOR into the lipid membrane. Successful incorporation of κOR onto the microtoroid sensor was confirmed by measuring the resonant wavelength shift and with fluorescence imaging (Fig. 3, Fig. 4b, Fig. S4). κOR-DynA 1-13 dose-response and competitive binding experiments were conducted entirely in Tris-HCl 25 mM, NaCl 150 mM, EDTA 1mM, BSA 0.05 % (w/v), pH 7.5. For radioligand binding^70^, cell membranes were incubated in 12×75 mm glass tubes with 5 nM ^3^H-naloxone with varying concentrations of DynA 1-13 at 25°C for 30 min. Receptor bound radiolabel was separated from free radiolabel by dilution and vacuum filtration over Whatman GF/C filters. Filters were counted in a liquid scintillation counter.

### Constructing Binding Curves

The binding curves were constructed by plotting the peak value vs. their respective ligand concentration. The dissociation constant, K_d_, was determined by fitting the peak values with the Hill-Waud model, which takes into account binding cooperativity and multiple binding sites by introducing the Hill coefficient, *n*^52,53^:

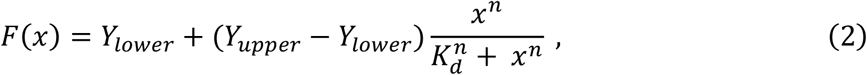

where *Y*_*lower*_ is the lower limit (fm), *Y*_*upper*_ is the upper limit (fm), *x* is the ligand concentration (M), *K*_*d*_ is the dissociation constant (M), and *n* is the Hill coefficient of cooperativity.

To account for experimental variations, the peak values for each experiment were normalized using the upper and lower limits from the fit curve:

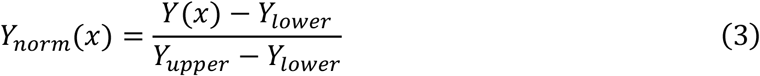

Where *Y(x)* is the peak value to be normalized as a function of concentration *x, Y*_*lower*_ is the lower limit of the fit curve (fm), and *Y*_*upper*_ is the upper limit of the fit curve (fm).

The inhibition constant, K_i_, of naloxone was calculated from the IC_50_ by fitting the competitive binding curve with a two-site competitive binding model:

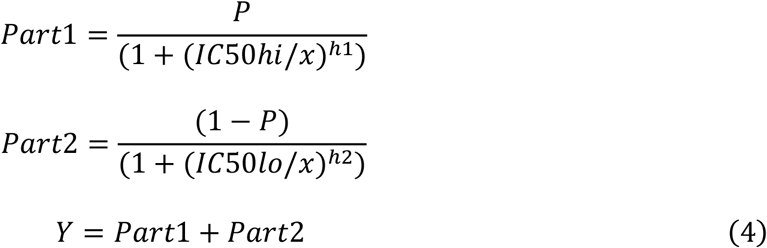

Where *Top* and *Bottom* is the upper and lower limit, respectively, of the fit curve. *P* is the fraction of high affinity binding sites for the competitor. *IC50hi* and *IC50lo* are the high and low IC50 values, respectively. *h1* and *h2* are the Hill slopes for the high and low IC50 values.

and the K_i_ was calculated from the IC50 using the equation:^71^

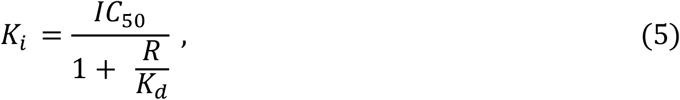

where *R* is the fixed Dynorphin A concentration (10 nM). *K*_*d*_ is characterized in Fig. 4d and is 3.1 nM.

The limit of detection (LOD) was determined as the lowest concentration analyte likely to be discernable from the buffer sample and was estimated using the equation:^72^

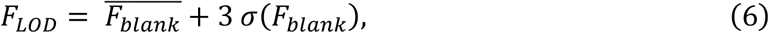

Where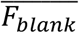 is the mean signal from blank samples and σ(F_blank_) is the standard deviation of the signal from 3 blank samples. The blank values were determined by taking the peak value from the wavelength shift of triplicate buffer injections in the LOD experiment (Fig. 5). The LOD is then the concentration where the calibration response curve intercepts FF_LoD_ in Fig. 5b.

### Data Acquisition

The microtoroid resonance frequency was tracked using the top-of-fringe locking function on a Toptica Digilock 110. The Digilock voltage output was connected to the laser’s frequency modulation input and an analog voltage data acquisition card (DAQ) (National Instruments PCI-4461, Austin, TX, USA). The Digilock modulates the laser’s frequency with a small amplitude 2 kHz sine wave and generates an error signal. As the microtoroid’s resonance frequency changes, the Digilock compensates by sending a voltage signal to the laser’s frequency modulation input which is also recorded by the DAQ card.

### Data Processing

Typical sources of noise (e.g., electrical noise at 60 Hz) are filtered out in the Fourier domain. Finally, a median filter is used to smooth the data. The parameters from doseresponse curves, competitive binding curves and statistical analysis were calculated using OriginPro 2023. Error bars show the standard error from 3-4 different experiments.

### COMSOL Multiphysics Simulation

A two-dimensional axisymmetric COMSOL simulation was used to compute the interaction between the microtoroid dielectric surface (refractive index *n* = 1.45) and the functionalized GM1-DOPC lipid bilayer (*n* = 1.42, total thickness [lipid + hydration layer] = ~9 nm) in buffer solution (*n* = 1.3385). The circular cross-section of toroid has a major radius of 45um and a minor radius of 2um. The value of the azimuthal mode number m of Equation (1) was set to 535 corresponding to a wavelength of 778.8nm. The field distribution and Q-factor of the WGM were analyzed using an eigenfrequency solver at a minimal mesh size of 0.4nm. An imaginary component (10^-7^) of the cavity refractive index was introduced to represent the scattering, absorption and other losses to lower the Q-factor to the actual experimental value. It is clearly seen that a portion of the optical field extends into the buffer sensing region.

