## Supplemental Information for "Label-free, real-time monitoring of membrane binding events at zeptomolar concentrations using frequency-locked optical microresonators"

**Address correspondence to:

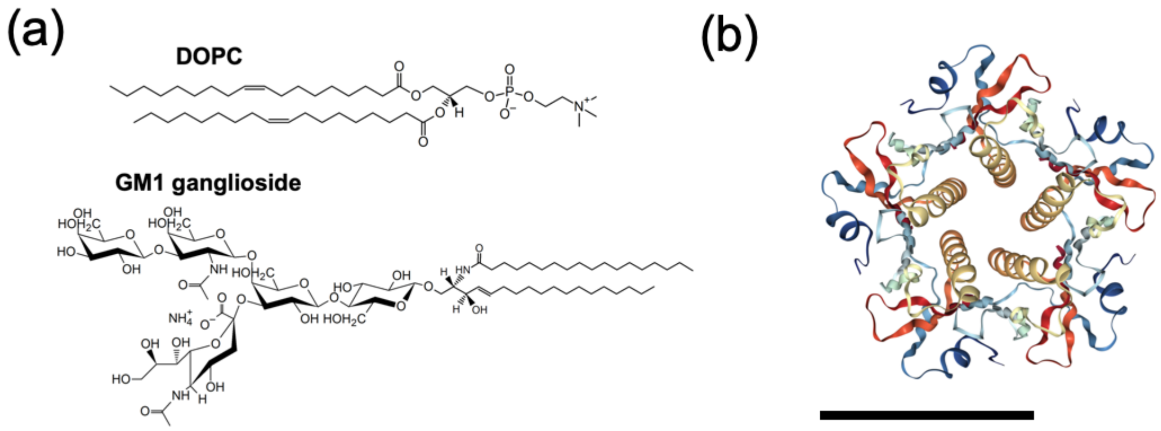


**Fig. S1** Chemical structure of the DOPC lipid and GM1 receptor glycolipid investigated in this study. (b) Pentameric structure of the cholera toxin B-subunit (Image from RCSB PDB (rcsb.org) of 1FGB (Zhang, R.G., Westbrook, M.L., Westbrook, E.M., Scott, D.L., Otwinowski, Z., Maulik, P.R., Reed, R.A., Shipley, G.G. (1995). The 2.4 Å crystal structure of cholera toxin B subunit pentamer: choleragenoid. J.Mol.Biol. 251: 550-562). The scale bar is 3.6 nm.

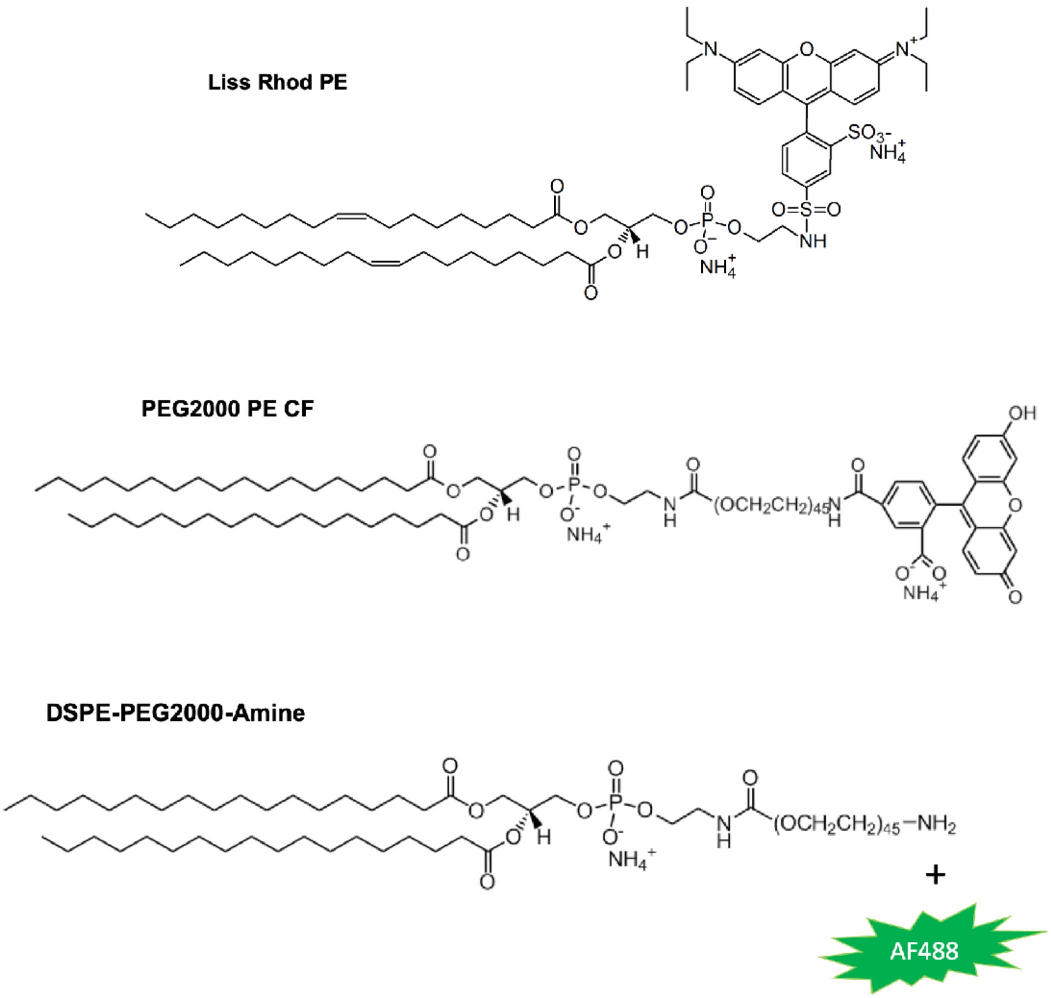


**Fig. S2**. Chemical structures of fluorescently labeled lipids used in this study. For the FRAP experiments, DSPE-PEG2000-Amine labeled with AF488 NHS ester was used to achieve better photostability than 18:1 PE CF (1,2-dioleoyl-sn-glycero-3-phosphoethanolamine-N-(carboxyfluorescein)).

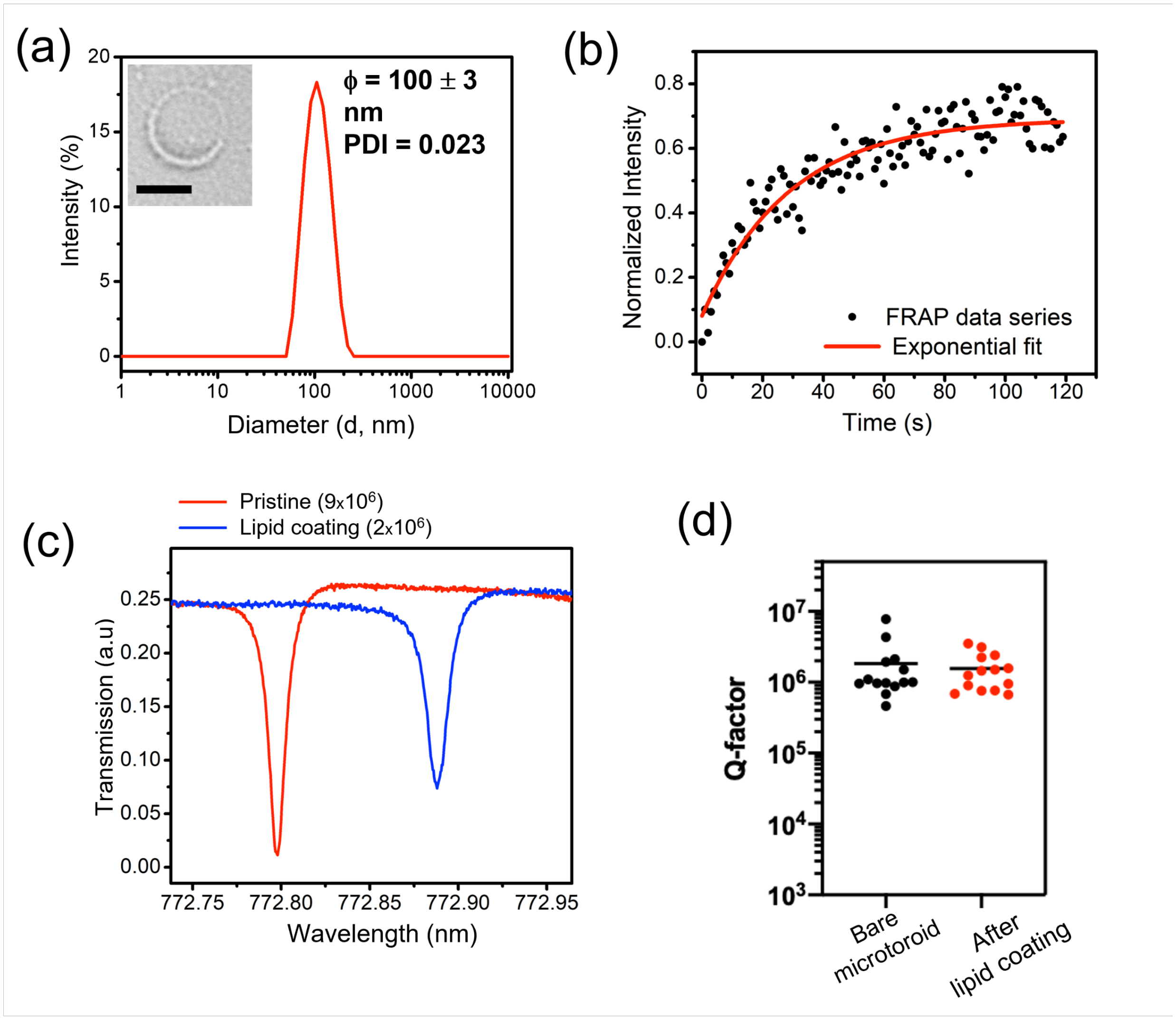


**Fig. S3.** Coating microtoroids with GM1-DOPC lipids. (a) Dynamic light scattering (DLS) measurement of GM1-DOPC lipid vesicles, showing a narrow size distribution centered at 100 nm. The inset picture is a TEM image of GM-DOPC lipid vesicles revealed by uranyl acetate staining (scale bar 50 nm). (b) FRAP experiment showing the fluorescence recovery of the lipid supported microtoroid. (c, d) Quality (Q)-factor measurement of microtoroid optical resonators before and after lipid coating (all the measurements were done in aqueous condition, recorded from different microtoroids on different days of experiments (n=14).

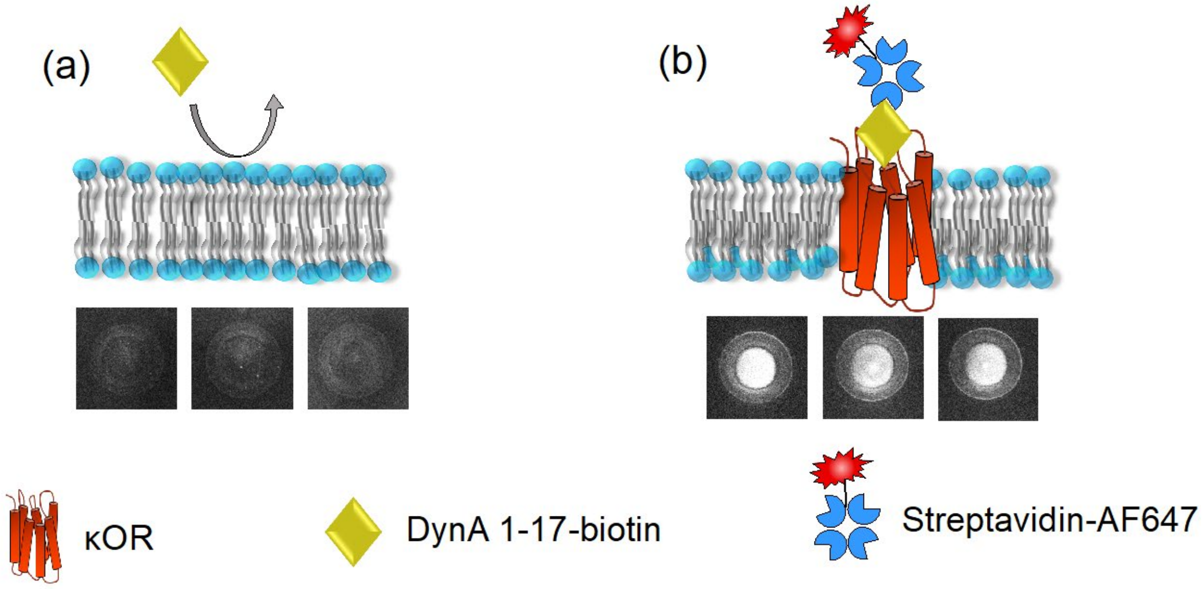


**Fig. S4.** Fluorescent imaging experiment to confirm the specificity of biotin DynA 1-17 to KOR reconstituted lipid coated microtoroid sensors. The fluorescent label used is streptavidin-AF647. (a) Control experiment of DynA 1-17 binding to a KOR-free, lipid-coated microtoroid sensor. As expected, little binding is observed. (b) DynA 1-17-streptavidin binding to a κOR reconstituted lipid coated microtoroid sensor. A bright fluorescence was revealed.

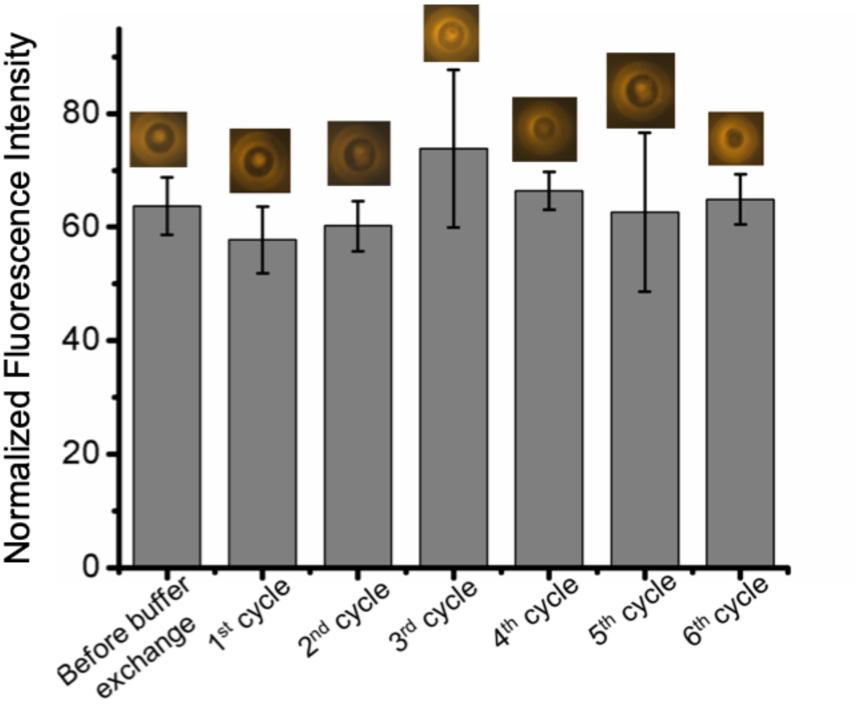


**Fig. S5**. Lipid functionalization stability during several cycles of complete buffer exchange (Tris 25 mM, NaCl 150 mM, pH 7.4). Liss Rhod PE doped DOPC lipid was used in this experiment. The average fluorescent intensity was taken from 3 random microtoroids on one chip. The fluorescence was normalized to the background intensity.

**Investigation of GM1-CTB toroid surface regeneration.** To achieve a high-throughput study for measuring GM1-CTB interactions using the microtoroid sensor, the CTB saturated surface was regenerated using a regeneration solution known to disrupt the interaction of CTB to GM1 receptors (i.e. 6M Urea, 25 mM Glycine, 160 mM NaCl, pH 2.5).[71] The regeneration efficiency was confirmed by a fluorescent imaging assay. CTB labeled with AF647 was bound to a GM1-DOPC membrane labeled with carboxyfluorescein (PEG2000 PE CF). The decrease in AF647 after regeneration directly related to the restored GM1 binding sites. As expected, after introducing the regeneration solution, the fluorescence intensity from CTB-AF647 significantly decreased while the fluorescence from the lipid membrane remained (Fig.S5). After 10 cycles of regeneration (by complete buffer exchange, Fig.S5), the CTB binding capacity still sufficiently remained. The regeneration efficiency was not significantly different between 2 consecutive cycles (2 min per cycle). The optimal regeneration time was 2 minutes. Each regeneration cycle was followed by binding buffer wash supplemented with 0.5 % BSA to prevent non-specific binding which may result from unexpected lipid removal.

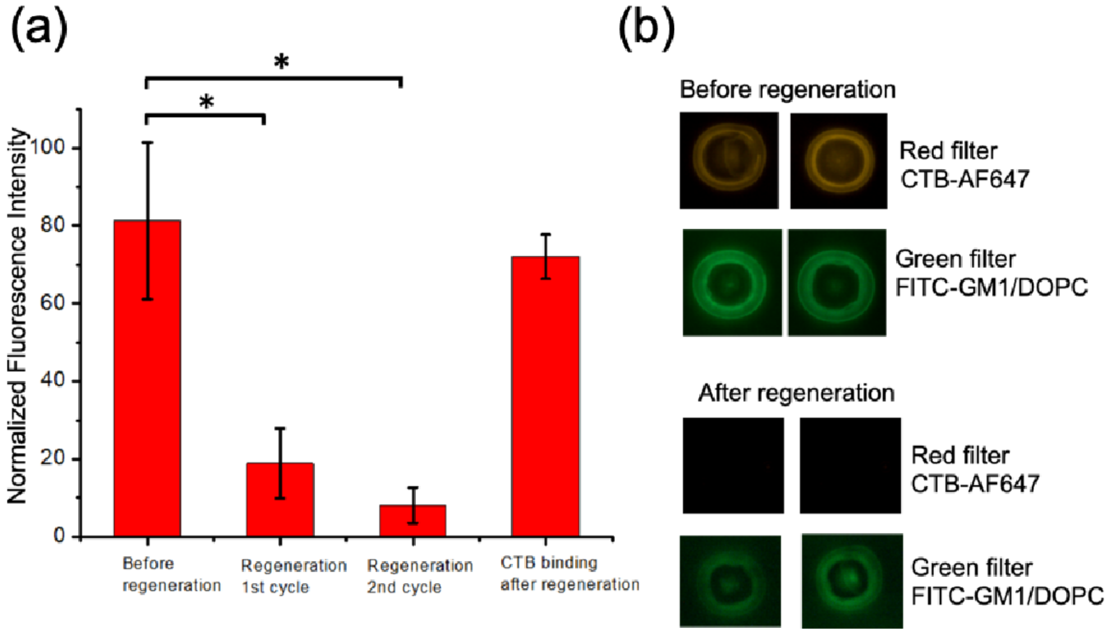


**Fig. S6.** CTB-GM1 regeneration using regeneration buffer (glycine 25 mM, Urea 6 M, NaCl 0.1 M, pH 3) characterized by fluorescent imaging (a) Plot of normalized fluorescent intensity of CTB-AF647 binding to a GM1-DOPC coated microtoroid before and after a regeneration step (statistical evidence $*p < 0.05$ performed by one-way ANOVA, n = 6) No significant difference in regeneration efficiency was observed between the two washes (2 mins each). The GM1-DOPC binding efficacy is high even after 10 regeneration cycles on-chip, (b) Corresponding fluorescent intensity images of CTB-AF647 binding to a GM1-DOPC supported microtoroid surface. The red fluorescence is from CTB-AF647 and the green fluorescence is from FITC labeled GM1-DOPC. Fluorescence imaging was performed for visualization of the retained lipid membrane after regeneration (FITC labeled GM1-DOPC intensity data was normalized with the images taken in the binding buffer (Tris 25 mM, NaCl 150 mM) only).

**Direct deposition of KOR membrane vs. KOR reconstitution.** The KOR/DynA pharmacokinetic study requires the surface of the silica toroid sensor to be functionalized with the receptors embedded in a lipid membrane. HEK293T+KOR cells were suspended in buffer and mechanically disrupted with a syringe and needle. We initially tried incubating the toroid sensors in a solution of crude HEK293T cell membrane fragments containing kOR directly on the toroid. However, the toroid Q-factor was degraded significantly because natural cell membranes contain many other additional proteins besides our protein of interest as well as larger cell fragments contributing to increased surface roughness and scattering loss. To maintain high Q factors, we instead remove the kOR from the crude cell membrane by detergent solubilization and reconstitute it in an artificial DOPC lipid bilayer.[1] This has the added benefit of dispersing the GPCRs, which can be tightly packed in a natural membrane, restricting ligand access to binding sites. Therefore, our approach *extends,* rather than restricts, general applicability. In general, an engineered membrane-receptor system has the potential for wider applicability than a natural-receptor system. For our other reported experiments (cholera toxin B binding to GM1), we obtained the GM1 in purified form and did not need to extract it from a natural membrane, and so there was no Q factor degradation in those experiments either.

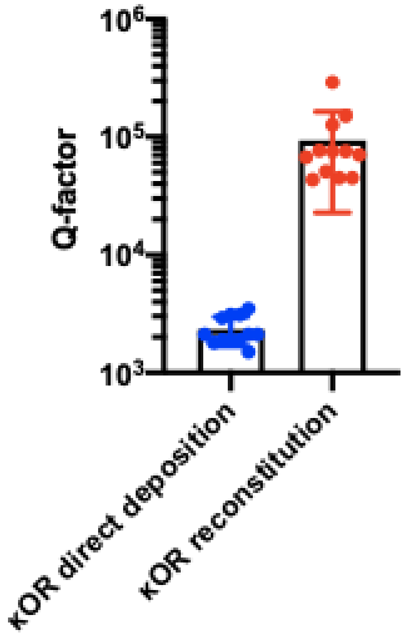


**Fig. S7.** Quality (Q)-factor data showing Q degradation when using direct deposition of a crude cell membrane vs reconstitution of kOR in a lipid bilayer ($n$ = 12). Direct deposition of kOR-proteolipid membrane degrades the microtoroid quality factor because the cell membrane contains many other additional proteins besides our protein of interest. For our sensor system, we achieve higher quality factors if we remove the kOR from the crude cell membrane and reconstitute it in a lipid bilayer. Tightly packed GPCRs restrict access to binding sites[2]; this is avoided if we reconstitute the proteins in a lipid bilayer.

**Experimental Setup.**

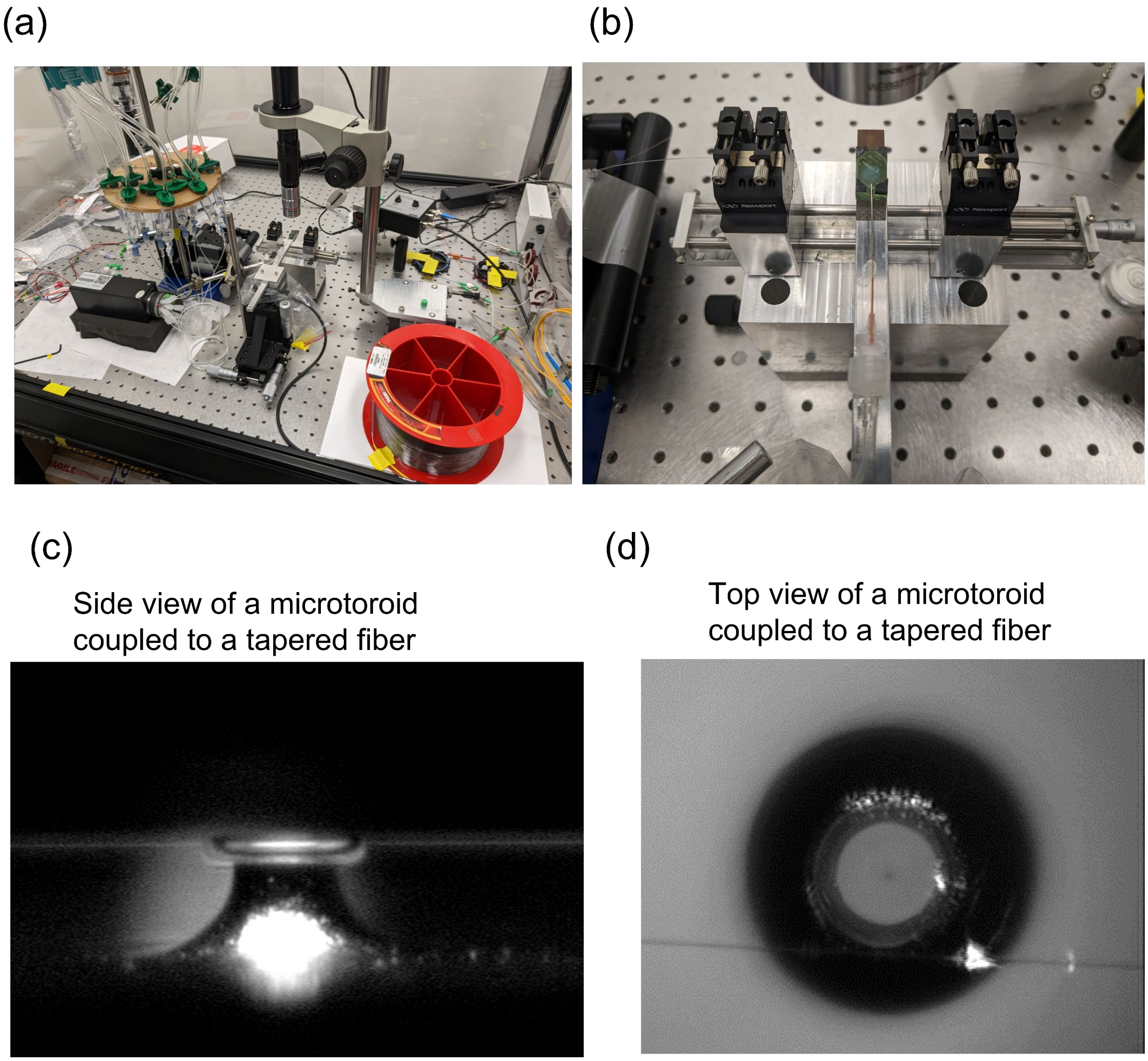


**Fig. S8.** Experimental setup and view of microtoroid and tapered fiber. (a) Experimental setup of FLOWER system: pressurized perfusion system and rotary valve for delivering fluid samples (left). Top-view microscope, fluidic chamber, tapered fiber stage, and piezoelectric nano-positioning cube on a 3-axis micrometer stage (middle). Single-mode fiber spool, in-line polarizer, attenuator, and photodetector (right). (b) Closer view of the fluidic chamber (green) positioned in the tapered fiber stage, along with the perfusion needle which delivers liquid samples into the fluidic chamber. (c) Side view of microtoroid and tapered fiber. (c) Top view of the microtoroid coupled to a tapered fiber.

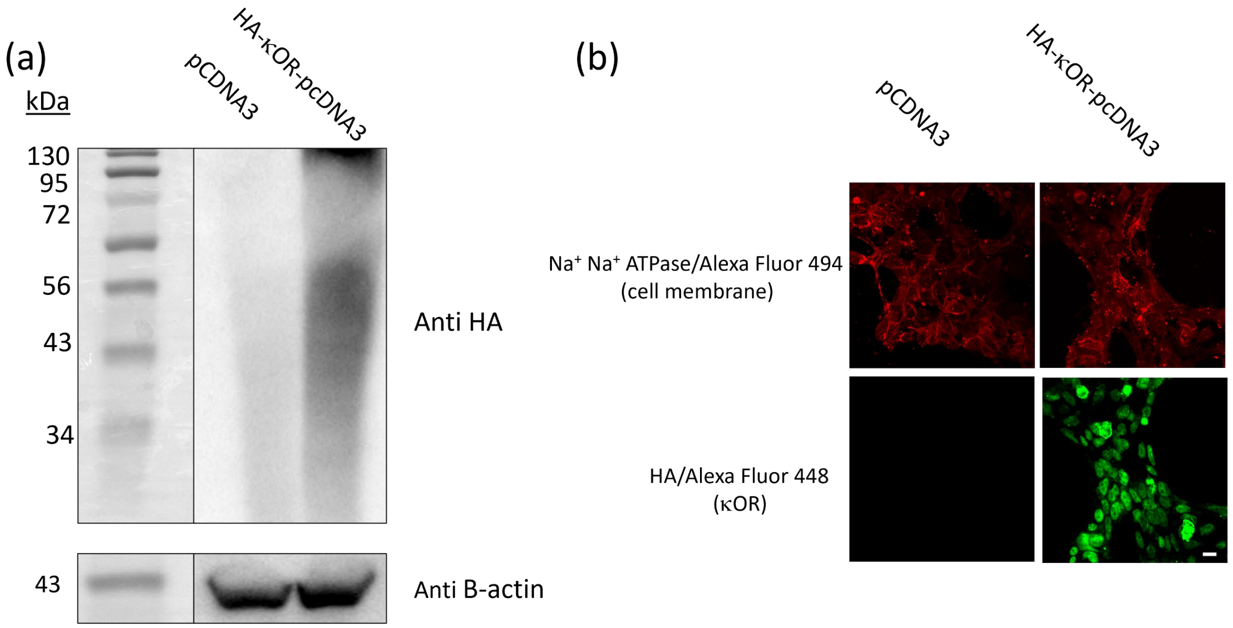


**Fig. S9.** Expression of kOR in HEK293T cells. A) Immunoblot of cell lysates showing expression of kOR only from the cells transfected with the HA-kOR-pcDNA3 expression vector. B) Confocal microscopy of nonpermeabilized HEK-293T transfected with empty vector or the kor expression vector. Red = cell membrane from incubation with Na+ K+ ATPase conjugated to Alexa Fluor 594. Green = expression of the HA tagged kOR from incubation with HA antibody conjugated to Alexa Fluor 448. Expression was detected in cells transfected with the HA-tagged kOR but not the empty vector control. 80X zoom; bar = 50 mm. Results are shown for representative experiments of 3 performed.
